# Plasticity of the *MFS1* promoter leads to multi drug resistance in the wheat pathogen *Zymoseptoria tritici*

**DOI:** 10.1101/174722

**Authors:** Selim Omrane, Colette Audéon, Amandine Ignace, Clémentine Duplaix, Lamia Aouini, Gert Kema, Anne-Sophie Walker, Sabine Fillinger

## Abstract

The ascomycete *Zymoseptoria tritici* is the causal agent of septoria leaf blotch on wheat. Disease control relies mainly on resistant wheat cultivars and on fungicide applications. The fungus displays a high potential to circumvent both methods. Resistance against all unisite fungicides has been observed over decades. A different type of resistance has emerged among wild populations with multi-drug resistant (MDR) strains. Active fungicide efflux through overexpression of the major-facilitator gene, *MFS1,* explains this emerging resistance mechanism. In this study, we identified as responsible mutations three types of inserts in the *MFS1* promoter, two of which harboring potential transcription factor binding sites. We show, that type I insertion leads to *MFS1* overexpression and consequently to MDR. Interestingly, all three inserts correspond to repeated elements of the *Z. tritici* genome.

These results underline the plasticity of repeated elements leading to fungicide resistance in *Z. tritici* and which contribute to its adaptive potential.

## Introduction

Wheat is the most widely grown crop in the world. It is subject to several diseases, principally due to fungal pests. Its major disease in Europe and North-America is septoria leaf blotch (SLB) caused by *Zymoseptoria tritici* (formerly *Mycosphaerella* g*raminicola*) (Fones & Gurr, 2015, Torriani *et al.*, 2015). The disease pressure of SLB depends on epidemical and environmental factors (Suffert *et al.*, 2011) and can be reduced by adapted agronomical practices (e.g., crop rotation) and intelligent use of less susceptible varieties (Brown *et al.*, 2015). Finally, SLB prevention strongly relies on the application of fungicides, namely inhibitors of sterol demethylation (DMIs, including azoles), inhibitors of mitochondrial complex II (SDHIs) and the multisite inhibitor chlorothalonil. According to disease pressure, spray programs targeting SLB range from one (Southern Europe) to four (Ireland and UK) sprays and around two treatments/year in France. The first treatment generally includes mixtures of azoles and chlorothalonil. The second spray aims to protect the key stage when the first leaf is emerging. Mixtures of azoles and SDHIs are often applied.

*Z. tritici* populations have developed resistance to all unisite fungicides, but to different extends. Azole resistance is generalized in Europe since the 1990s and now affects field efficacy of the molecules, but SDHI resistance has just emerged and does not entail yet the efficacy of this mode of action (Lucas *et al.*, 2015). Resistance is principally due to target site modification or overexpression (Cools & Fraaije, 2013, Fraaije *et al.*, 2011, Leroux & Walker, 2011).

Multi drug resistance (MDR) operating through increased drug efflux is a resistance mechanism recently detected in some field isolates of *Z. tritici*. Since it is associated to azole target site resistance, it confers high resistance factors to this class of inhibitors, whereas only low resistance levels towards SDHIs are recorded (Leroux & Walker, 2011).

The phenomenon of MDR is well known from human cancer cells and antibiotic resistant bacteria (Wu *et al.*, 2014, Hiramatsu *et al.*, 2014). In fungi, *Saccharomyces cerevisiae* has served as model organism to elucidate MDR (also termed PDR for “pleiotropic drug resistance”) and its regulation. It has also been extensively studied in several pathogenic yeast species, e.g., *Candida albicans* and *C. glabrata*. For reviews we refer the reader to some excellent papers (Gulshan & Moye-Rowley, 2007, Moye-Rowley, 2003, Paul & Moye-Rowley, 2014, Cannon *et al.*, 2009, Morschhäuser, 2010). Globally, MDR is conferred to by constitutive overexpression of membrane transporter genes either of the ATP-binding cassette (ABC) type or of the major facilitator superfamily (MFS). These transporters expulse drugs outside the cell, thereby reducing the intracellular drug concentrations. Their specificity can be more or less broad (Higgins, 2007). Constitutive overexpression of membrane transporters in clinical isolates of *C. albicans* was found to be due to gain-of-function mutations in the transcription factors Tac1 or Mrr1, controlling respectively the expression of the ABC transporter gene *CDR1* or of the MFS protein encoding *MDR1* gene.

In the phytopathogenic fungi, *Botrytis cinerea, Sclerotinia homeocarpa* and *Oliculimacula yallundae* MDR has also been described for field isolates (Chapeland *et al.*, 1999, Leroux *et al.*, 1999, Hulvey *et al.*, 2012, Sang *et al.*, 2015, Leroux *et al.*, 2013). The mutations responsible for MDR in *B. cinerea* field strains have been identified. They correspond either to a retro-element like insert in the promoter of the *BcmfsM2* gene or to gain of function mutations in the transcription factor Mrr1 controlling the expression of the ABC transporter gene *BcatrB* (Kretschmer *et al.*, 2009, Leroch *et al.*, 2013). Strains harboring either or both mutations are frequent among *B. cinerea* wild populations (Walker *et al.*, 2013, Leroch *et al.*, 2013, Rupp *et al.*, 2016).

In a recent study we have shown that fungicide efflux was at work in two *Z. tritici* MDR field isolates (Omrane *et al.*, 2015). Both strains, as well as other MDR strains tested, constitutively overexpress the gene *MFS1* (originally named *MgMFS1* for *Mycosphaerella graminicola MFS1*) encoding an MFS transporter, capable to transport a wide diversity of molecules (Roohparvar *et al.*, 2008). Its inactivation in one MDR strain abolished the MDR phenotype revealing that the MFS1 protein is necessary for the MDR phenotype at least in this strain. In both analyzed MDR strains, a 519 bp insert was detected in the *MFS1* promoter, a putative reminiscence of an ancient LTR-retrotransposon. Other, but not all field MDR strains, proved to have this promoter insert as well, suggesting a potential role in MDR (Omrane *et al.*, 2015). In this study we address the question of the mutations responsible for the MDR phenotype in the previously characterized MDR strains.

We used Bulk Segregant Analysis (BSA) in order to map mutations responsible for MDR in *Z. tritici*. BSA is a genotyping method adapted to monogenic traits (Michelmore *et al.*, 1991). It is based on the establishment of two phenotypically dissimilar pools derived from an offspring population. These pools are genotyped and markers linked to the phenotype according to their allelic frequencies are selected to determine the locus of interest. With the rise of Next Generation Sequencing (NGS), statistical tools have been developed for BSA phenotyping to uncover QTLs (Magwene *et al.*, 2011, Duitama *et al.*, 2014) and others (Liu *et al.*, 2012). BSA has been applied to identify genomic *loci* contributing to natural polymorphism (Leeuwen *et al.*, 2012) or mutant phenotypes (Duveau *et al.*, 2014). In fungi the combination of BSA, was successfully used to identify mutations responsible for cell cycle and developmental processes respectively in *Neurospora crassa* (Pomraning *et al.*, 2011) and *Sordaria macrospora* (Nowrousian *et al.*, 2012). Due to its haploid, characterized and annotated genome (Goodwin *et al.*, 2011) and the possibility to perform sexual crosses between different strains (Kema *et al.*, 1996), *Z. tritici* is suitable to BSA to map the mutation(s) responsible for the MDR phenotypes.

In this work we mapped the mutation responsible for the MDR phenotype in the two previously characterized field strains using BSA. After functional validation of the responsible mutation, the 519 bp promoter insert of the *MFS1* gene, we screened *Z. tritici* field strains for the *MFS1* promoter genotype. Interestingly two other inserts were identified in the same promoter only in *Z. tritici* MDR field strains overexpressing *MFS1*.

## Materials & Methods

### Z. tritici strains and growth conditions

All *Z. tritici* field strains used in this study are listed in Table 1. As sensitive reference strains, we used IPO323 and IPO94269 displaying *MAT1-1* and *MAT1-2* genotypes respectively (Waalwijk *et al.*, 2002), and sensitive to all tested fungicides (Roohparvar *et al.*, 2007 and our unpublished results). As principal MDR strains we used 09-ASA-3apz and 09-CB01 (Omrane *et al.*, 2015, Leroux & Walker, 2011). Strains produced in this study are described below. All strains (if not otherwise indicated) were cultivated on solid YPD medium (20 g L^-1^ Bacteriological peptone, 10 g L^-1^ Yeast extract, 20 g L^-1^ Glucose, 15 g L^-1^ agar) during 7 days at 17°C in the dark. The sequence of IPO323 is publically available (Goodwin *et al.*, 2011), that of IPO94269 was kindly provided by Syngenta AG for mapping analysis.

**Table 1:**
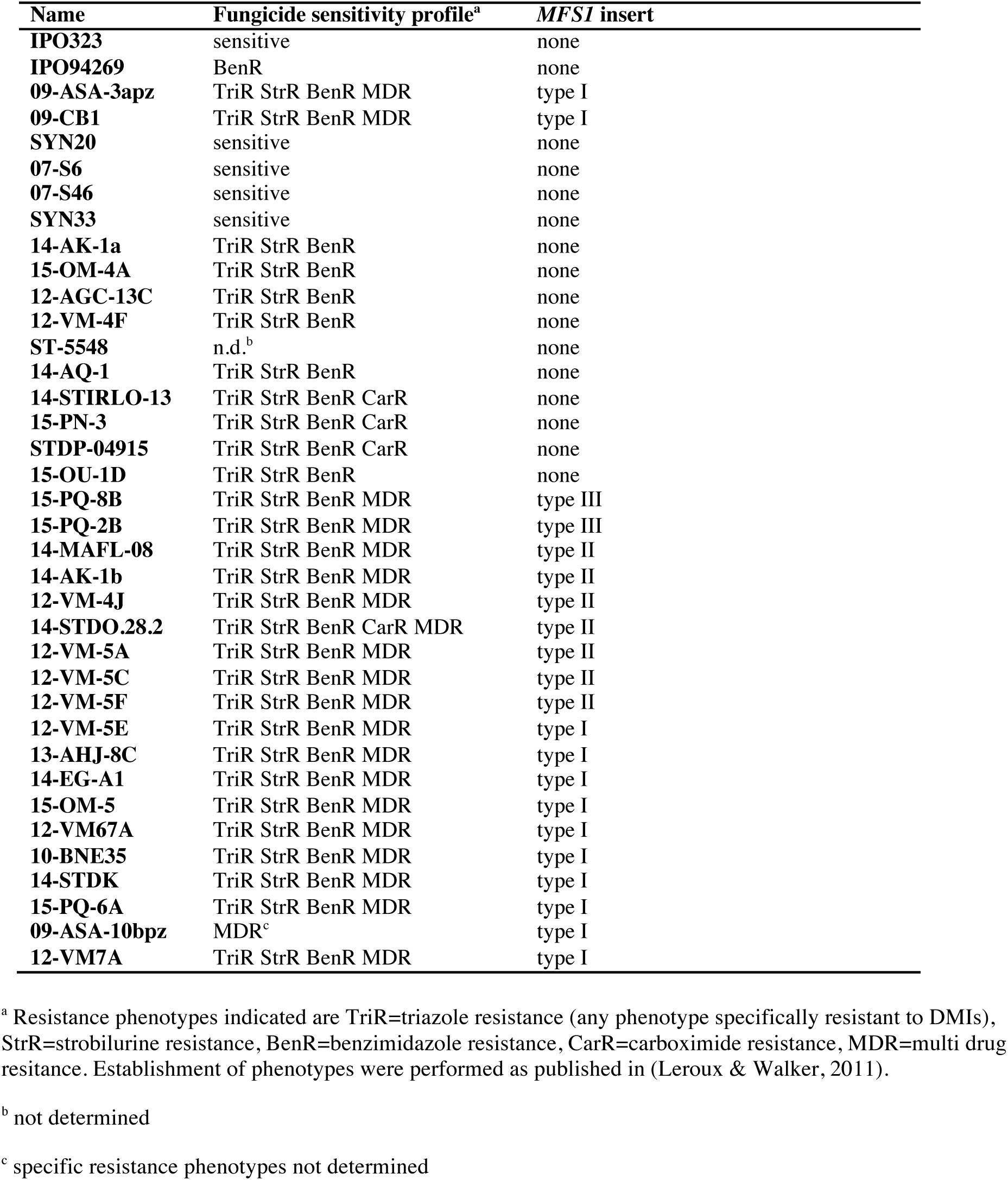
*Zymoseptoria tritici* field strains used in this study.

To determine EC_50_ values to prothioconazole-desthio, tebuconazole, metconazole, tolnaftate, terbinafine and pyrifenox, conidia were collected from 3 day-old cultures on NY medium in sterile water and adjusted to a final concentration of approximately 2x10^5^ conidia mL^-1^. 300μL of each solution was spread on test plates containing solid phosphate-glucose medium (2 g L^-1^ K_2_ HPO_4_; 2 g L^-1^ K_2_H PO_4_; 10 g L^-1^ glucose, 12.5 g L^-1^) with 5 fold serial dilutions (2.5 fold) of prothioconazole-desthio (Sigma-Aldrich, Saint Quentin Fallavier, France) tebuconazole (technical grade, BayerCropScience, Germany), metconazole (technical grade, BASF Agro, Germany), tolnaftate (Sigma-Aldrich), terbinafine (technical grade, Sandoz, Switzerland) and pyrifenox (technical grade, Syngenta Agro, Switzerland) as described by (Leroux & Walker, 2011). All fungicides were supplied as 250x concentrated ethanol-solutions. Test and control plates were incubated at 17°C in the dark for 48 h. The length of germ-tube was estimated microscopically on 10-30 spores per plate. The EC_50_ value of each tested fungicide corresponding to the concentration inhibiting spore germination by 50% was determined by non-linear regression (least squared curve fitting) using the GraphPad PRISM program, (GraphPad software, La Jolla, CA, USA).

### Crosses and progeny phenotyping

Crosses 1 (09-ASA-3apz x 09-CB1), 2 (09-ASA-3apz x IPO94269) and 3 (09-CB1 x IPO323) were performed as described previously (Kema *et al.*, 1996). Single spore progenies were isolated from ascii. A set of 20 progeny strains from each cross were checked by genotyping of 11 SSRs on the core chromosomes (Gautier *et al.*, 2014) in comparison to their parents to validate the absence of external contaminants prior to all subsequent analyses. The whole offspring was genotyped using a set of eight SSRs. Only strains presenting single bands were conserved.

Sensitivity tests to distinguish MDR from sensitive offspring were performed in solid as well as in liquid YPD medium with the addition of tolnaftate (2 μg mL^-1^, 5 μg mL^-1^or 10 μg mL^-1^). All strains were grown on solid YPD for one week (18°C; continuous light) and transferred by toothpicks to a 96-well microtiter-plate into 200 μL sterile water. 10 μL of 1/100 dilutions was spotted on 96 microtiter plates filled with YPD (liquid) with and without tolnaftate. The plates were incubated on a rotary shaker at 18°C during 11 days. Scoring of growth was made by O.D. measurement (λ= 590 and 620 nm) at 3, 6 and 11 days. Growth rates at days 6 and 11 were calculated relative to day 3 and as a ratio of treated *vs*. untreated conditions. The whole assessment procedure was repeated three times.

### Progeny bulk preparation

Resistant (MDR phenotypes) and sensitive sets (sensitive phenotypes) of strains were selected on the basis of the phenotyping described above among those that grew or not on 5 μg mL^-1^ of tolnaftate to build bulks for each of the progenies (crosses 2 and 3) before nucleic acid extraction. The number of strains for the DNA bulks was respectively n = 60 for R2 and S2 (resistant and sensitive bulks, respectively from cross 2), n = 50 for R3 and S3 (resistant and sensitive bulks, respectively from cross 3). The phenotype of all selected strains was verified on tolnaftate (5 μg mL^-1^) by growth tests on solid and liquid medium. Additionally, liquid growth tests were performed with or without the addition of reversal agents namely amitriptyline, chlorpromazine and verapamil (Sigma-Aldrich) at a 1:3 ratio (5 μg mL^-1^tolnaftate, 15 μg mL^-1^reversal agents).

For DNA extraction each strain was grown individually in liquid YPD during 1 week on a rotary shaker (140 rpm; 18°C). Cultures were harvested by centrifugation (4000 rpm during 10 min.), rinsed twice with cold PBS 1X (Sigma-Aldrich, Saint-Quentin, France) and immediately frozen in liquid nitrogen prior to vacuum-freeze-drying. DNA were extracted from freeze dried cells using the DNeasy Plant maxi kit (Qiagen) for. The concentrations were determined and the quality verified on a 2100 Bioanalyzer (Agilent). Identical amounts (200 ng) of genomic DNA of each strain were mixed to constitute the bulks R2, S2, R3 and S3.

### Sequencing and mapping to reference genome

100 base paired-end Illumina sequencing was performed on 2 μg of DNA of each bulk as well as for the MDR progenitors by the sequencing provider (Beckman-Coulter). The total number of reads used for genome mapping was 32269.4 Mb distributed equally among the DNA samples (Table S1). These were mapped respectively on reference genome sequences of the sensitive parental strains IPO94269 (reads from 09-ASA-3apz and R2, S2 bulks) or IPO323 (09-CB1 and R3, S3 bulks) by the sequencing provider, using Bwa 0.6.1 (Li & Durbin, 2009) for alignment and the GATK GenomeAnalysis module (McKenna *et al.*, 2010) for local realignment. Variants were called with Samtools 1.18 (Li *et al.*, 2009) using default parameters. The GATK Unified Genotyper (McKenna *et al.*, 2010) was used to call all locations of allele frequencies. For 09-CB1 and R3, S3 bulks, snpEff 3.0 was used to call effects for filtered variants. Finally, VarSifter 1.5 (Teer *et al.*, 2012) was used to inspect final genotype calls for coherence. IGV 2.3 (Robinson *et al.*, 2011) was used to visualize the polymorphisms (SNPs and INDELs) per cross project respectively. For cross 2 only contigs of IPO94269 over 200bp were used as reference sequences. For cross 3 the JGI Mycgr3 genome (http://genome.jgi.doe.gov/Mycgr3/Mycgr3.home.html) was used together with the “FrozenGeneCatalog20080910” gene predictions.

Alignments of both reference sequences were performed by MAUVE (Darling *et al.*, 2004). Reads’ quality cutoff and read depth were respectively set to ≥ 20 and ≥ 5. Final filtering parameters to detect the genomic distortions were set as GQ scores (encoded as a phred quality) for each genotype as ≥ 0.5 for resistant bulks and < 0.5 for sensitive bulks, while the parental genotypes were set as = 1 (MDR) and 0 (Sensitive reference sequence). The difference between R and S bulk genotypes (D value) was filtered as ≥ 0.4. Blastn of contig sequences were made at http://genome.jgi-psf.org/pages/blast.jsf?db=Mycgr3 using *M. graminicola* v2.0 unmasked nuclear assembly as search criteria.

### Genotyping of progeny strains

#### MFS1 promoter

Screening for the 519 bp insert in the *MgMFS1* promoter (Omrane *et al.,* 2015) among all progeny strains was performed by PCR using the primer couple MFS1_2F and MFS1_4R (Table S2). No promoter insert lead to a 486 bp amplicon while the insert increased the amplicon size to 1005 bp.

#### NFX1, PYC polymorphism

The genotypes of the *NFX1* and *PYC* genes located at both ends of contig 1135 of the IPO94269 genome sequence were determined on the progeny strains by HRM in comparison to the parental strains. The primer couples NFX1_11422FW / NFX1_11422RV and PYC_5UTR_FW / PYC_5UTR_RV (Table S2) showing close to 100% amplification efficiency on serial DNA dilutions of the parental strains were used for HRM comparisons under the following conditions. In a total volume of 25 μL, 5 ng of genomic DNA was analyzed with 300 nM of both primers using SsoFast(tm) Evagreen^®^ Supermix (Bio Rad, Marnes-la-Coquette) at 1x. The cycling parameters were 98°C for 2 min followed by 40 cycles of 98°C for 2 sec and 60°C for 5 sec. The high resolution melt curve was established between 70°C and 95°C with 0.2°C increments every 10 sec in a CFX-384 Real-Time PCR System (Bio Rad). Melting curve normalization and differentiation was performed using Precision Melt Analysis Software (Bio Rad).

#### MFS1 expression analysis

For *MFS1* expression during exponential growth, the tested strains (field strains, offspring, transformants) were grown in liquid YPD (5 mL) at 18°C and 140 rpm during 48-72 h. The cell concentration was determined microscopically with a hematocytometer and diluted in 100 mL of YPD to a final concentration of 5x10^4^ cells/mL and incubated for 48 h at 18°C, 140 rpm to a final cell concentration of 5x10^6^ cells/mL. The cultures were harvested by centrifugation (6500 rpm, 4°C during 20 min), the pellet immediately frozen in liquid nitrogen and freeze-dried.

Total RNA was extracted from the freeze-dried mycelium using the RNeasy Plant Mini kit (Qiagen) according to the supplied instructions. Quality of the RNA was checked by electrophoresis, the concentration determined spectrophotometrically (NanoDrop(tm), ThermoScientific(tm)). 1 μg of total RNA was used for cDNA synthesis using the PrimeScript(tm) RT Reagent Kit with gDNA Eraser (Takara). cDNAs were diluted 5 times before qPCR analysis with MESA GREEN qPCR MasterMix Plus for SYBR^®^ Assay (Eurogentec, Angers, France). The MFS1 amplicon was obtained with primers 110044_Fw and 110044_RV which had been verified by standard curves for amplification efficiencies ranging from 95% to 105 % and for the absence of non-specific amplicons. Relative expression levels were determined according to using the 2^-ΔC(t)^ and the bestkeeper method (Pfaffl *et al.*, 2004) with *EF1α, β-tubulin* and *UBC1* as control genes to establish the most stable value of housekeeping gene expression among all experimental conditions. Means and standard deviations were calculated from two technical replicates of two biological replicates. All primer sequences are listed in Table S2.

#### MFS1 gene replacement constructs

To introduce the various *MFS1*^*MDR*^ alleles into the sensitive IPO323 strain, the following replacement cassettes were constructed. The respective *MFS1* allele, 1380 bp upstream until 518 bp downstream of the ORF, was amplified from the corresponding DNA (09-ASA-3apz, 09-CB01 or other MDR strains) using the primers MDR-pKr_F at the 5’ end and the strain specific primer MDR6_hygR (09-ASA-3apz) or MDR7_hyg R (09-CB01) at the 3’ end using Q5^®^ High-Fidelity DNA Polymerase (New England Biolabs, Evry, France). A 737 bp 3’ flank of the MFS1 gene to facilitate homologous recombination was amplified from IPO323 genomic DNA with primers Ipo323-hygroF and Ipo323-pKraR. Finally the hyromycine resistance marker *hph* was amplified from plasmid pCAMB-HPT-Hind (Kramer *et al.*, 2009) with the primer couples Hygro_MDR6_F / Hygro_ipo323_R or Hygro_MDR7_F / Hygro_ipo323_R respectively. The three fragments (0.06 pmol of each) were assembled with *Xho*I-*Eco*RI digested pCAMB-HPT-Hind (0.02 pmol) by Gibson Assembly^®^ Cloning Kit (New England Biolabs, Evry, France) according to the supplier’s instructions. Half of the assembly reaction was used to transform NEB 5-alpha competent *Escherichia coli* (New England Biolabs). The kanamycine resistant colonies were PCR-screened with the above-cited primers. Positve clones were picked for plasmid extractions according to standard protocols (Green & Sambrook, 2012). The extracted plasmids were checked again by *Eco*RI-*Nco*I restriction.

The resulting plasmids pCAMBIA-MFS1(MDR6) and pCAMBIA-MFS1(MDR7) respectively were introduced into *Agrobacterium tumefaciens* AGL1 competent cells by heat-shock. Transformants were selected and isolated on LB broth with rifampicin (20 μg.mL^-1^), kanamycin (50 μg.mL^-1^) and ampicillin (50 μg.mL^-1^).

#### Z. tritici transformation and analysis

Agrobacterium mediated transformation procedure was performed as described by Zwiers and de Waard, (2001) according to the modifications made by Kramer et al., (2009) with minor changes. Transformants were selected and isolated on hygromycin (100 μg.mL^-1^) containing YPD medium. 50 transformants for each plasmid were picked and isolated twice on selective YPD medium. All purified transformants were tested by PCR: genomic DNA was extracted from cells harvested on solid YPD medium with the GenElute^(tm)^ Plant Genomic DNA Miniprep Kit (Aigma-Aldrich, Saint-Quentin Fallavier, France). Z4_110044_FW and Z4_110044_RV primers (Table S2) were used to PCR amplify the *MFS1* promoter in order to distinguish transformants devoid of the insert (700 bp amplicon) from those with the insert (1200 bp amplicon) and from ectopic integrations (700 bp with additional amplicon). 16 and 10 transformants respectively out of 50 (20-32%) had the *MFS1*^*MDR*^ allele integrated at the *MFS1 locus* of IPO323.

#### Fungicide sensitivity assays of selected transformants

All validated transformants were tested for their sensitivity to various fungicides on solid YPD medium supplemented with tolnaftate (2 μg mL^-1^), terbinafine (0.015 μg mL^-1^), prochloraz (0.05 μg mL^-1^), metconazole (0.02 μg mL^-1^), boscalid (2 μg mL^-1^), bixafen (0.5 μg mL^-1^) and azoxystrobin (20 μg mL^-1^). All fungicides were supplied as 1000x concentrated ethanol-solutions. Strains were precultered in 5 mL of liquid YPD medium during 3 days at 18°C and 140 rpm. After OD_600_ measurement, all cultures were adjusted with fresh YPD medium to the lowest measured OD. Three sequential 10-fold dilutions were prepared from all adjusted precultures. 3 μL of precultures and dilutions were spotted on each fungicide assay and control plate and incubated at 17°C in the dark during 5 days.

## Results

### I. Genetic mapping of *mdr loci* in two MDR field strains

To check whether MDR phenotypes described in (Leroux & Walker, 2011) are driven by allelic mutations, we performed a cross between strains 09-ASA-3apz and 09-CB1. Rapid discrimination of MDR strains from sensitive ones consists in a growth test using fungicides that are not used in agriculture such as tolnaftate and terbinafine (Leroux *et al.*, 1999), both squalene epoxidase inhibitors, typically used against human fungal infections (Barrett-Bee *et al.*, 1986, Georgopapadakou & Bertasso, 1992). A progeny of 140 strains was analyzed on tolnaftate. Growth tests did not reveal any sensitive isolate (Table 2), suggesting that the two parental *mdr* mutations are closely linked on the same chromosome, in a common genomic region.

**Table 2:** Crosses used in this study. Progeny segregation into sensitive or MDR is based on sensitivity/resistance to tolnaftate 2 μg mL^-1^.

|  | Cross (n° / strains) | sensitive | MDR | Total |
| --- | --- | --- | --- | --- |
| 1 | 09-ASA-3apz (MDR) x 09-CB1 (MDR) | 0 | 140 | 140 |
| 2 | 09-ASA-3apz (MDR) x IPO94269 (sens.) | 149 | 148 | 297 |
| 3 | 09-CB1 (MDR) x IPO323 (sens.) | 111 | 97 | 208 |

Both MDR strains were then crossed to the sequenced sensitive strains IPO94269 and IPO323, respectively. These two crosses were essential to map the *mdr loci.* We isolated and phenotyped 297 and 208 progeny strains respectively. Crosses 2 and 3 generated respectively 50% and 47 % MDR progeny strains (Table 2). In both cases the ratio MDR *vs.* sensitive strains was in agreement with a single mutation responsible for the MDR phenotype. However, we need to underline that the selection of progeny strains was not completely unbiased after the elimination of mixtures among progeny strains. Therefore, statistical tests cannot be applied to offspring segregation.

To map both *mdr* mutations, we decided to perform a bulk-progeny sequencing approach (BSA) as developed for different phenotypes in other fungal species (Nowrousian *et al.*, 2012, Pomraning *et al.*, 2011). MDR and sensitive progeny strains were selected on the basis of maximal phenotypic dissimilarities by growth tests in medium supplemented with tolnaftate and the membrane transporter inhibitor, verapamil to constitute resistant (R) or sensitive (S) bulks. We adjusted the number of progeny strains per bulk to n=50 (cross 3 09-CB1 x IPO323) and n=60 (cross 2 09-ASA-3apz x IPO94629). In order to map the *mdr* mutations, we adopted a 3-step protocol (Fig.1) involving (1) DNA extraction from each of the four bulks, (2) Illumina DNA sequencing and (3) sequence mapping to the reference sequence of the parental sensitive strain.

**Fig. 1:**
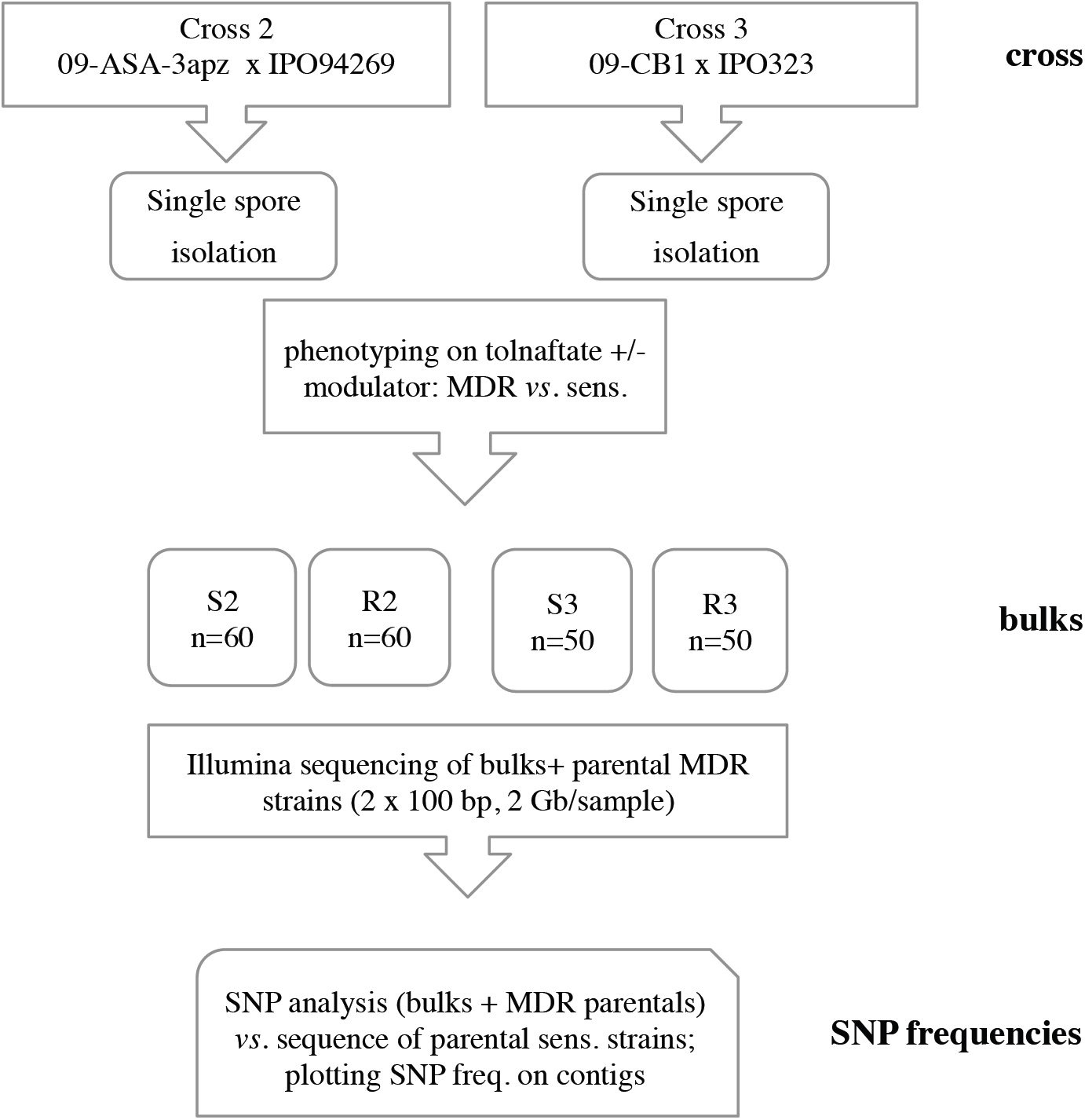
Flowchart of applied BSA procedure. The bulks of progeny strains are designated S and R as sensitive or resistant respectively to tolnaftate, the numbers 2 and 3 refer to the crosses listed in Table 2.

Reads derived from each bulk and from the MDR parental strains were mapped on their respective reference sequence IPO323 (IPO323, R3 & S3 reads) and IPO94269 (09-ASA-3apz, R2 & S2 reads). Mapping procedure yielded a comparable number of polymorphic sites (more than 180000) and density for both data sets (Table S1). SNP and INDEL frequencies relative to each reference sequence were calculated and reported as GQ-value between 0 and 1 (100%). We assumed that the GQ value of the region surrounding the *mdr* mutations would tend towards 1 in the resistant bulks, whereas the GQ values at the same sites would approach 0 in the sensitive bulks. By applying the thresholds GQ ≥ 0.5 to the resistant and GQ ≤ 0.5 to the sensitive bulks and considering as relevant only sites with a maximum difference between both bulks (GQ_R_-GQ_S_=D value ≥ 0.4) we were able to reduce the bin size around the distortion to several kilobases (Fig. 2B) located on the left arm of chromosome 7 of IPO323. In the 09-CB1 x IPO323 data set, this region showed the highest distortion, decreasing from the telomere to the centromere (Fig. 2B). The strategy was merely the same in the 09-ASA-3apz x IPO94269 dataset except for the use of unassembled contigs. 13 contigs out of 56 with D values > 0.4 co-localized on the left arm of chromosome 7 as well (Fig 2B), out of which contig 1135 had the highest number of polymorphic sites with the highest D values. We therefore focused our subsequent analysis on the region of chromosome 7 covered by this contig (from position 8 to 47 kb in Fig 2C). Reporting all SNPs and INDELS of the resistant bulks R2 and R3 present in this bin on the local alignment between contig 1135 and the IPO323 very left arm of chromosome 7, we identified three kinds of polymorphic sites: those common to both MDR strains and those independent to each of them (Table S3). The highest detected D values were respectively recorded in the *CYP52* gene (R2&R3; D value = 0.72; synonymous substitution), in the pyruvate carboxylate gene *PYC* (R2; D value = 0.738; intron) and the manganese-iron superoxide gene *Mn-SOD* (R3; D value = 0.736; 5’UTR). Intriguingly, this region also covers the transporter gene *MFS1*, whose involvement in drug efflux and MDR was shown before (Roohparvar *et al.*, 2007, Omrane *et al.*, 2015). This gene is located between the above-mentioned polymorphic sites. In particular, the genes immediately surrounding *MFS1, STK* and *PYC*, harbored many polymorphic sites that co-segregated with the MDR phenotype, while the 5’UTR and 3’UTR of the *MgMFS1* gene appeared structurally highly dissimilar in the 09-ASA-3apz and 09-CB1 backgrounds from both reference sequences as stated by the dramatic decrease of mapped reads (data not shown). Indeed, we had previously found that both strains harbor a 519 bp insert in the *MFS1* promoter region (Omrane *et al.*, 2015).

**Fig. 2:**
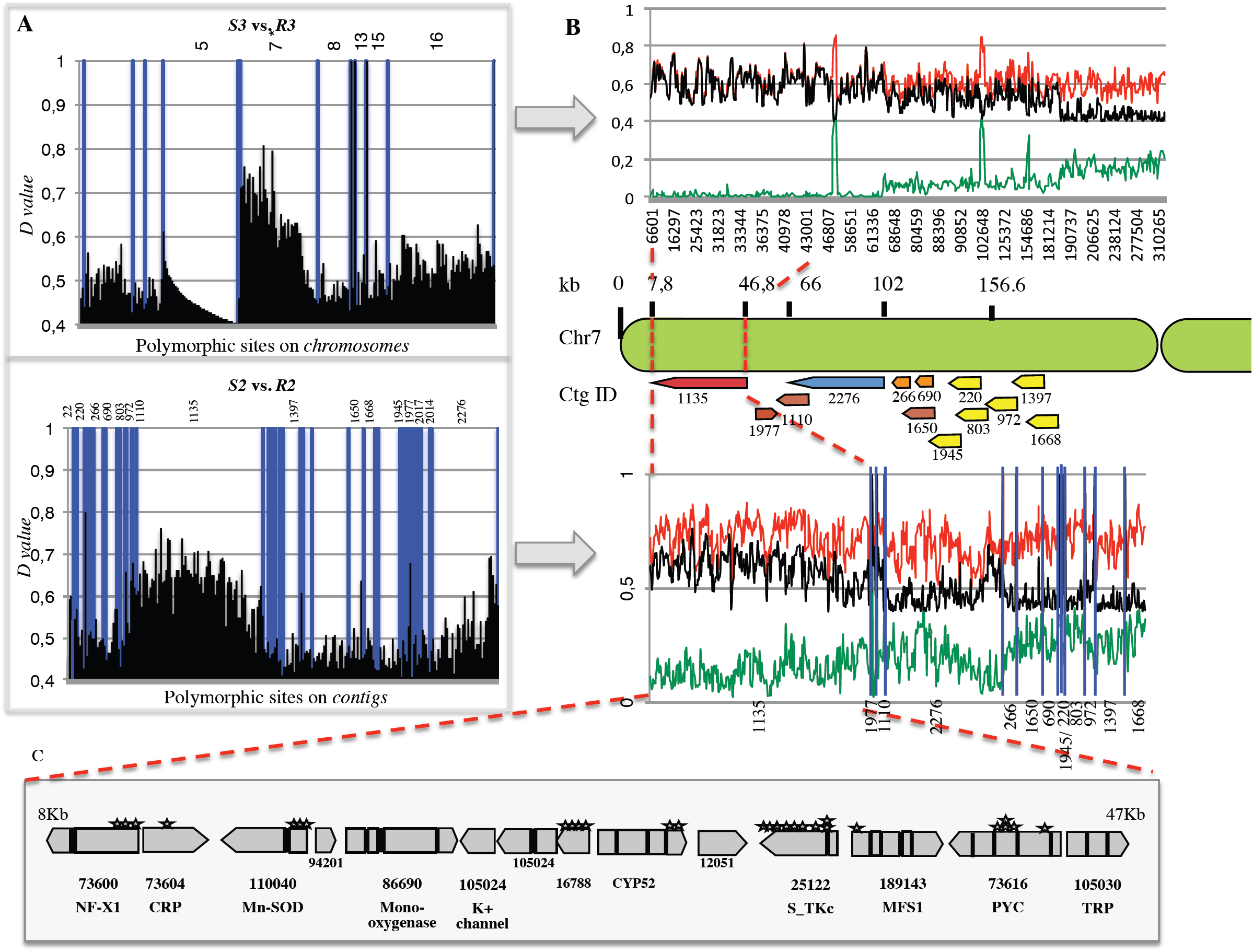
Polymorphism discovery in 09-CB1xIPO323 vs. 09-ASA-3apzxIPO94269 bulks. A/ D values (GQ_R_-GQ_S_) plotted against their chromosomes (IPO323=sensitive parent) or contigs (IPO94269=sensitive parent) derived from 09-CB1xIPO323 (upper panel) and 09-ASA-3apzxIPO94269 (lower panel). B/ Incrementing the bin size resolution of the regions concerned by the highest distortion, which shows a net increase of the D-values (black line) between resistant bulk’s (red lines) and sensitive bulk’s GQ scores. Contigs showing the highest distortion (containing at least one polymorphic site with D value ≥ 0.5) from the 09-ASA-3apzxIPO94269 bulk sequencing were selected and aligned against the IPO323 genome. C/ Contig 1135 matching the region extending from 8 to 47 kb of chromosome 7 with the highest D-values harbors 14 genes. SNPs and INDELs inducing non-synonymous substitutions in the coding regions are indicated by the stars.

Altogether, this BSA revealed for both MDR strains a region of 39 kb on chromosome 7 whose polymorphism strongly co-segregated with the MDR phenotypes. Since the cross between both MDR strains indicated that the responsible mutations are allelic or closely linked, the identification of this common region from both independent BSA experiments is in favor of its involvement in the MDR phenotype.

In order to precisely map the *mdr* mutation on the 39 kb fragment of 09-CB1, we analyzed the alleles of the marker genes at each extremity of the region by HRM, as well as the *MFS1* promoter genotype by PCR in all progeny strains derived from the cross between 09-CB1 and IPO323. Table 3 shows that only the *MFS1* promoter genotype of the MDR parental strain strictly co-segregated with the MDR phenotype. A 100% co-segregation between the *MFS1* promoter allele and the MDR phenotype was also observed for all 297 progeny strains derived from cross 2 between 09-ASA-3apz and IPO94269.

**Table 3:** Genotyping of progeny strains for *MFS1* alleles and linked marker genes.

| marker\genotype | MDR-progeny (n=97) |  | sensitive progeny (n=111) |  |
| --- | --- | --- | --- | --- |
|  | 09-CB1 | IPO323 | 09-CB1 | IPO323 |
| *NFX1* | 96 | 1 | 1 | 110 |
| *PYC* | 95 | 2 | 1 | 110 |
| *MFS1* | 97 | 0 | 0 | 111 |

| marker\genotype | MDR-progeny (n=148) |  | sensitive progeny (n=149) |  |
| --- | --- | --- | --- | --- |
|  | 09-ASA-3apz | IPO94269 | 09-ASA-3apz | IPO94269 |
| *NFX1* | n.a. | n.a. | n.a. | n.a. |
| *PYC* | n.a. | n.a. | n.a. | n.a. |
| *MFS1* | 148 | 0 | 0 | 149 |

Upper panel: cross 09-CB1 x IPO323; lower panel: cross 09-ASA-3apz x IPO94269; n.a. not analyzed

**Fig. 3:**
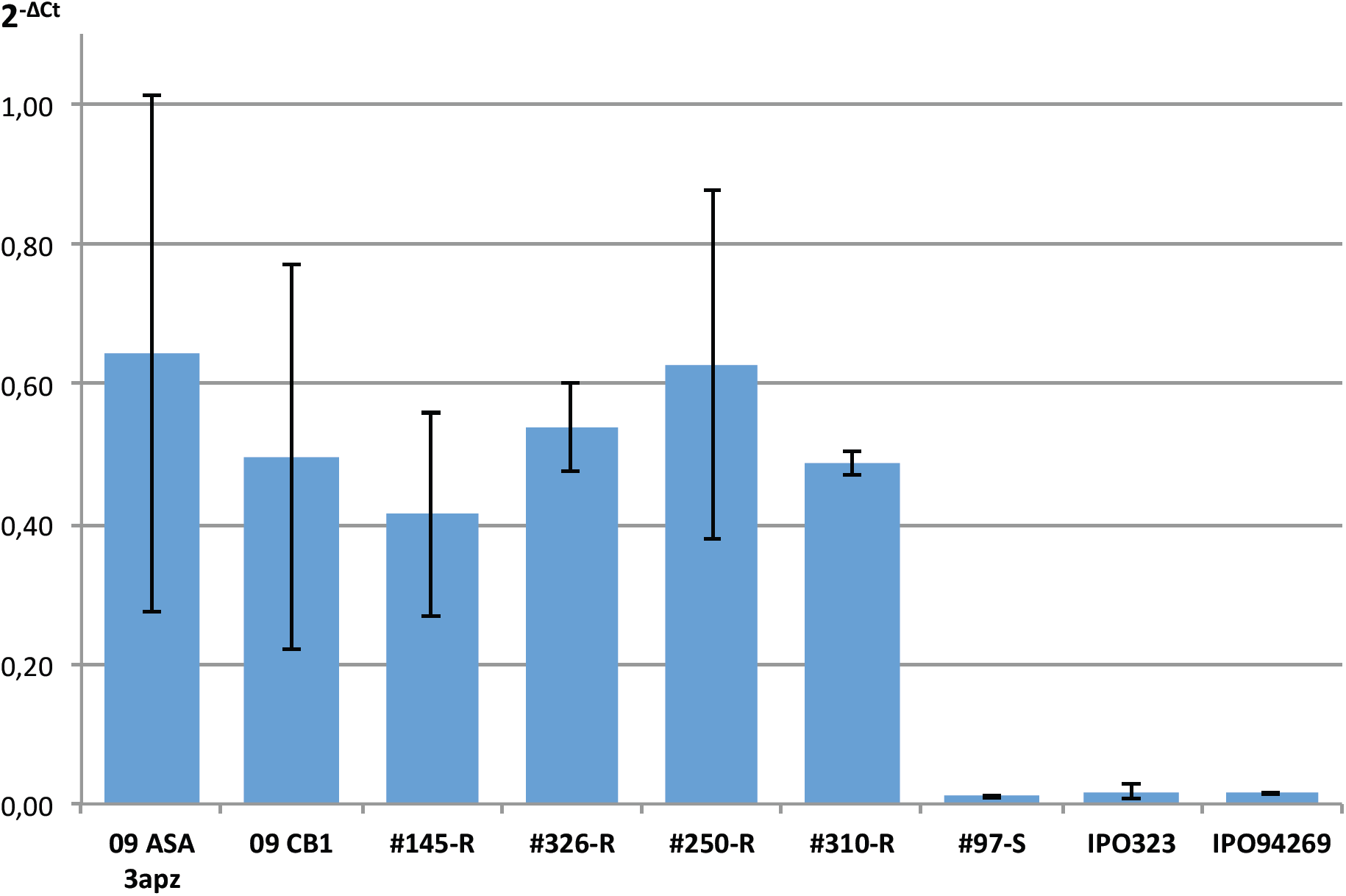
*MFS1* expression in parental and progeny strains. Values are means of two biological replicates, three or more in the case of the parental strains. Progeny strains with “R” suffix are of MDR phenotype. Strain #97-S is of sensitive phenotype.

Expression analysis of *MFS1* in selected offspring showed *MFS1* overexpression in resistant progeny strains (#145-R, #326-R, #250-R, #310-R) comparable to the MDR parental strains, while the sensitive strain #97-S displayed the basal expression level characteristic of sensitive strains (Fig. 3).

### Functional validation of the identified *mdr*mutation (*MFS1* insert)

In order to validate the involvement of the 519 bp promoter insert in the MDR phenotype, we proceeded through the replacement of the *MFS1*^*WT*^ allele by the *MFS1*^*MDR*^ allele in the sensitive reference strain IPO323 (Fig. 4 A, B). After selection and isolation on hygromycine, transformants were screened by PCR analysis with primers Z4_110044_FW and Z4_110044_RV (Table S2) flanking the promoter insert (Fig. 4 A, B), to discriminate those obtained by integration of the replacement construct at the *MFS1 locus* from those with ectopic integration, as latter showed more than one amplicon. Moreover, this PCR distinguished between integration events of type a and type b, *i.e.,* type a integration lead to the promoter insert, while type b integration did not (Fig. 4A).

**Fig 4:**
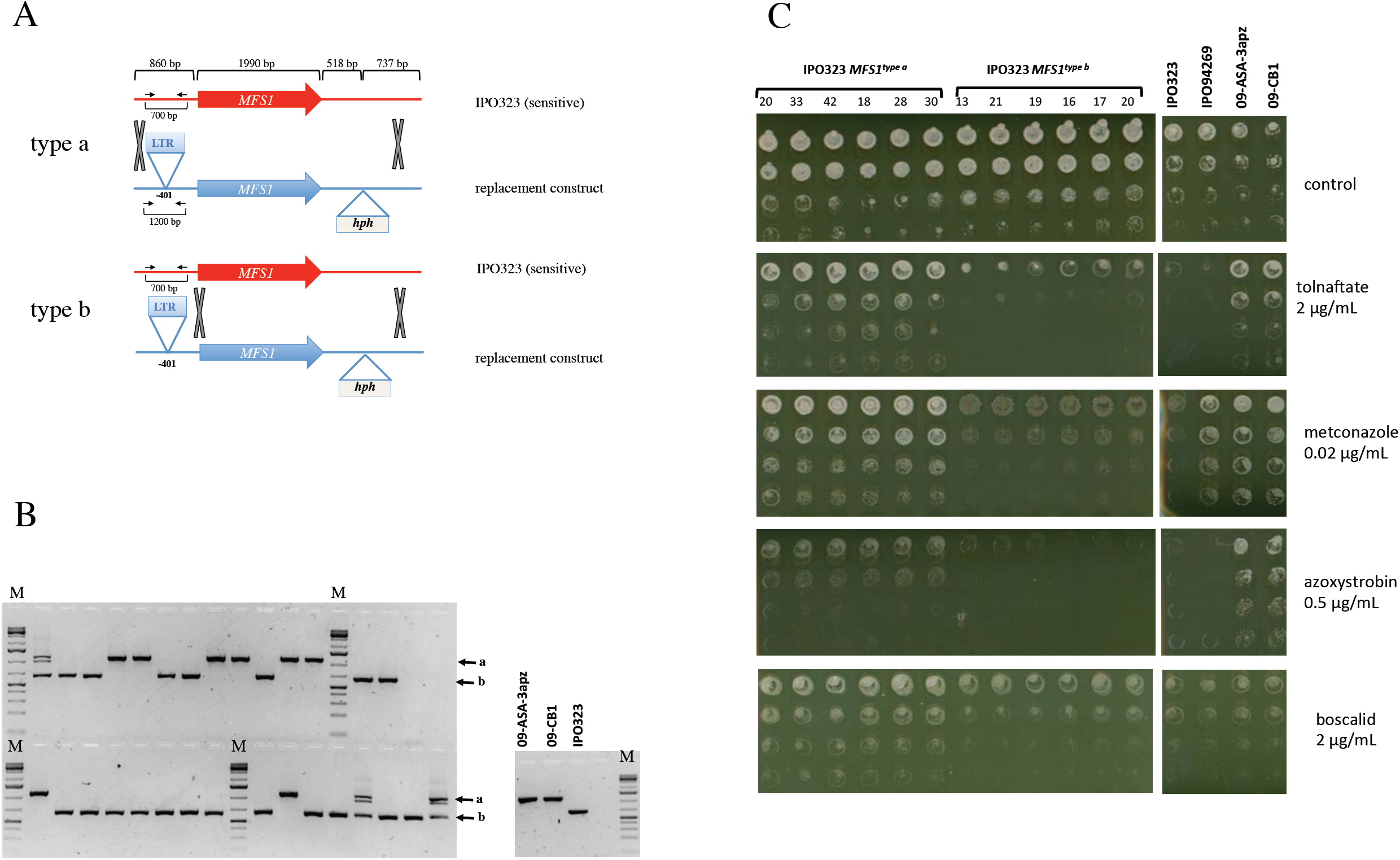
Functional analysis of the *MFS1*^*MDR*^ allele in the sensitive IPO323 strain. A/ Principle of possible homologous recombination events at the *MFS1 locus* with the replacement construct. The gray Xs indicate homologous recombination events leading to the integration of the hygromycine resistance marker *hph*. The black arrows indicate the positions of the primers 2F and 4R used to distinguish between type a and type b recombination events. B/ PCR analysis of isolated transformants with primers 2F and 4R. The black arrows indicate the bands produced by type a or type b recombination. C/ Growth tests of selected transformants on fungicides with different modes of action. Serial dilutions (top to down rows) of calibrated precultures of the indicated strains were inoculated on growth test plates (YPD with the indicated fungicides) and incubated at 17°C during 5 days.

Six transformants of each category were tested for growth on different fungicides. As visible from Fig. 4 C only the integration of the 519 bp insert in the *MFS1* promoter (type a integration) conferred increased tolerance to the squalene epoxidase inhibitor tolnaftate, to the DMI metconazole, to the QoI azoxystrobin and to the SDHI boscalid. Similar results were observed with terbinafine, prochloraz and bixafen (data not shown). Type b integrants displayed the same sensitivity to the compounds as the parental IPO323 strain. Four selected transformants with the promoter insert (transformants 20, 33, 18, 28) overexpressed *MFS1* compared to IPO323 while type b transformants 13, 19, 16, 17 (without the insert) did not (Fig. 5).

**Fig. 5:**
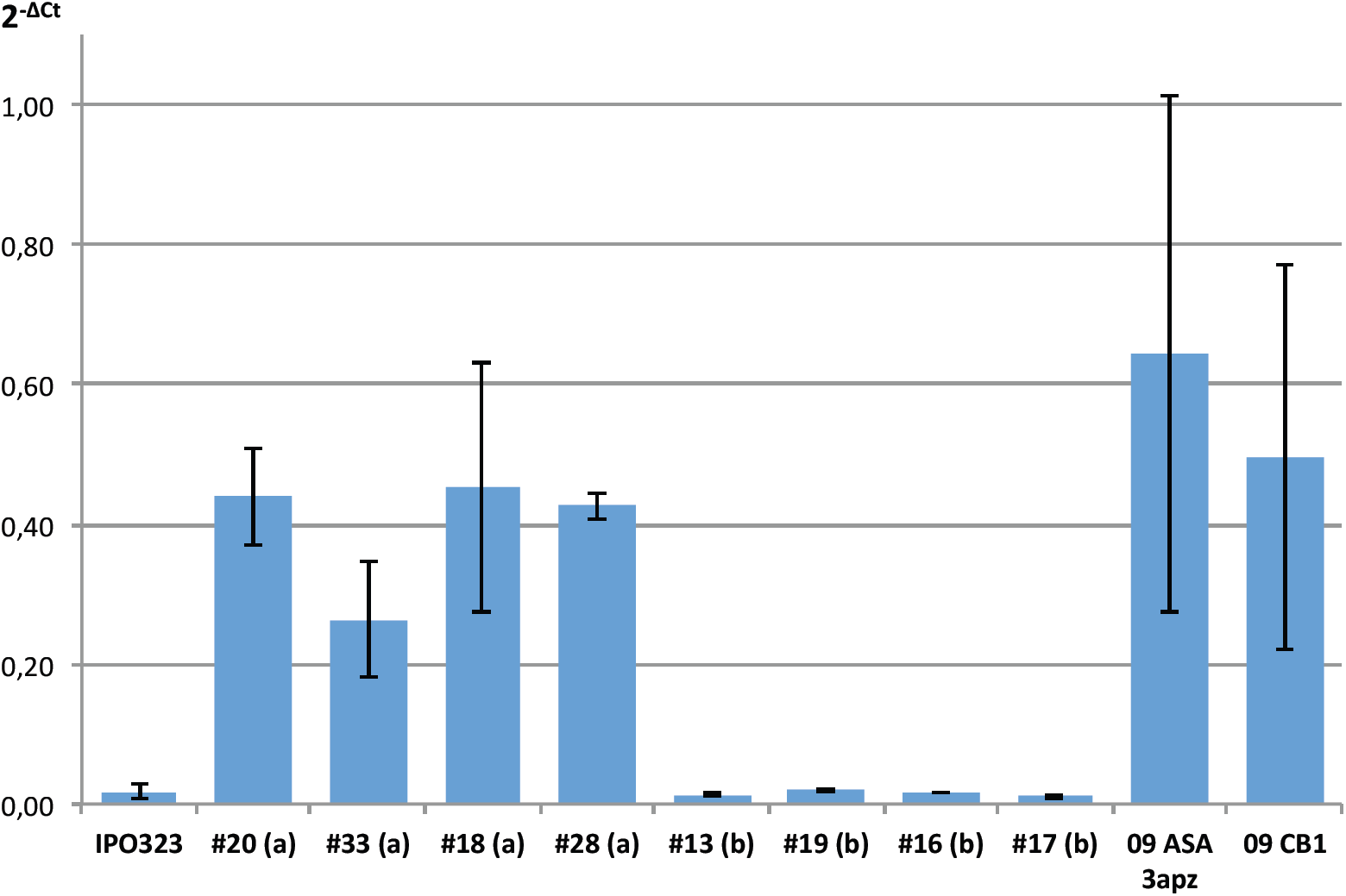
*MFS1*expression in IPO323 *MFS1*replacement mutants. *MFS1* expression was measured by qRT-PCR relative to three reference genes (*ß-tubulin, eF1a, actine)*. Values are means of two to three biological replicates, three or more in the case of the field strains. Transformants are indicated by the hashtag symbol (#). The letters in the brackets refer to the recombination event of Fig. 4. Type a recombination leads to the promoter insert of 519 bp and the MDR phenotype, while type b recombination event does not.

We may thus conclude that the 519 bp LTR insert in the *MFS1* promoter leads to its overexpression and consequently to an MDR phenotype.

### III.*MFS1*promoter and expression analysis in *Z. tritici* field isolates

In a population survey we tested if the *MFS1* promoter insert was present in all field isolates phenotyped as MDR, applying PCR with the primer pair MFS1_2F / MFS1_4R (Table S2). No insert was detected in non-MDR strains. In 35 genotyped MDR strains however, we obtained three different amplicons of 1000, 850 or 650 bp respectively, corresponding to three different inserts, designated type I, II or type III inserts. The 519 bp LTR insert (type I) was the most frequent insert, detected since 2009; type II insert was detected in 25% of the strains and present since 2012; type III insert was detected in only two strains from 2015 (Garnault et al., unpublished).

We determined the EC50 values on selected DMIs, terbinafine and tolnaftate for field strains representative each type of insert (n=7/type I; n=5/type II; n=2/type III). Resistance factors of the three genotypes to tolnaftate and terbinafine (unlinked to any specific resistance and not affected by the *CYP51* alleles), listed in Table 4, reveal similar phenotypes for strains with type I and II inserts, while the resistance factor conferred to by the type III insert seems weaker.

**Table 4:** Resistance factors (RF^a^) of *Z. tritici* field isolates with different *CYP51* and *MFS1* alleles relative to *CYP51*^*WT*^*MFS1*^*WT*^.

| genotype<sup>b</sup> |  | tolnaftate | terbinafine | epoxiconazole | prothioconazole-desthio | tebuconazole | metconazole | pyrifenox |
| --- | --- | --- | --- | --- | --- | --- | --- | --- |
| **CYP51<sup>WT</sup>**<br>**MFS1<sup>WT</sup>** | EC50<br>(ng mL<sup>-1</sup>) | 15 +/- 3.5 | 0.1 +/- 0.02 | 0.35 +/- 0.17 | 0.32 +/- 0.18 | 3.4 +/- 0.9 | 0.21 +/- 0.04 | 0.38 +/- 0.14 |
| **CYP51<sup>TriR</sup>**<br>**MFS1<sup>WT</sup>** | RF | n.d. | n.d. | 258 | 51.9 | 609 | 360 | 559 |
| **CYP51<sup>TriR</sup>**<br>**MFS1<sup>MDR-typeIII</sup>** | RF | 3.8 | 37.5 | 927 | 144 | 269 | 1116 | 709 |
| **CYP51<sup>TriR</sup>**<br>**MFS1<sup>MDR-typeII</sup>** | RF | 11.5 | 87.6 | 1138 | 107 | 381 | 572 | 1054 |
| **CYP51<sup>TriR</sup>**<br>**MFS1<sup>MDR-typeI</sup>** | RF | 11.9 | 84.3 | 959 | 73.6 | 546 | 655 | 610 |

<sup>a</sup> The RFs are expressed as EC50 (studied genotype) / EC50 (*CYP51*<sup>WT</sup> *MFS1*<sup>WT</sup>)
<sup>b</sup> The *CYP51*<sup>TriR</sup> genotype groups together different *CYP51* genotypes of field-strains with cross-resistance to DMIs

The *MFS1* transporter gene was found overexpressed also in the field strains with inserts of type II and type III (Fig. 6), although eventually at lower levels as compared to strains with insert of type I.

**Fig. 6:**
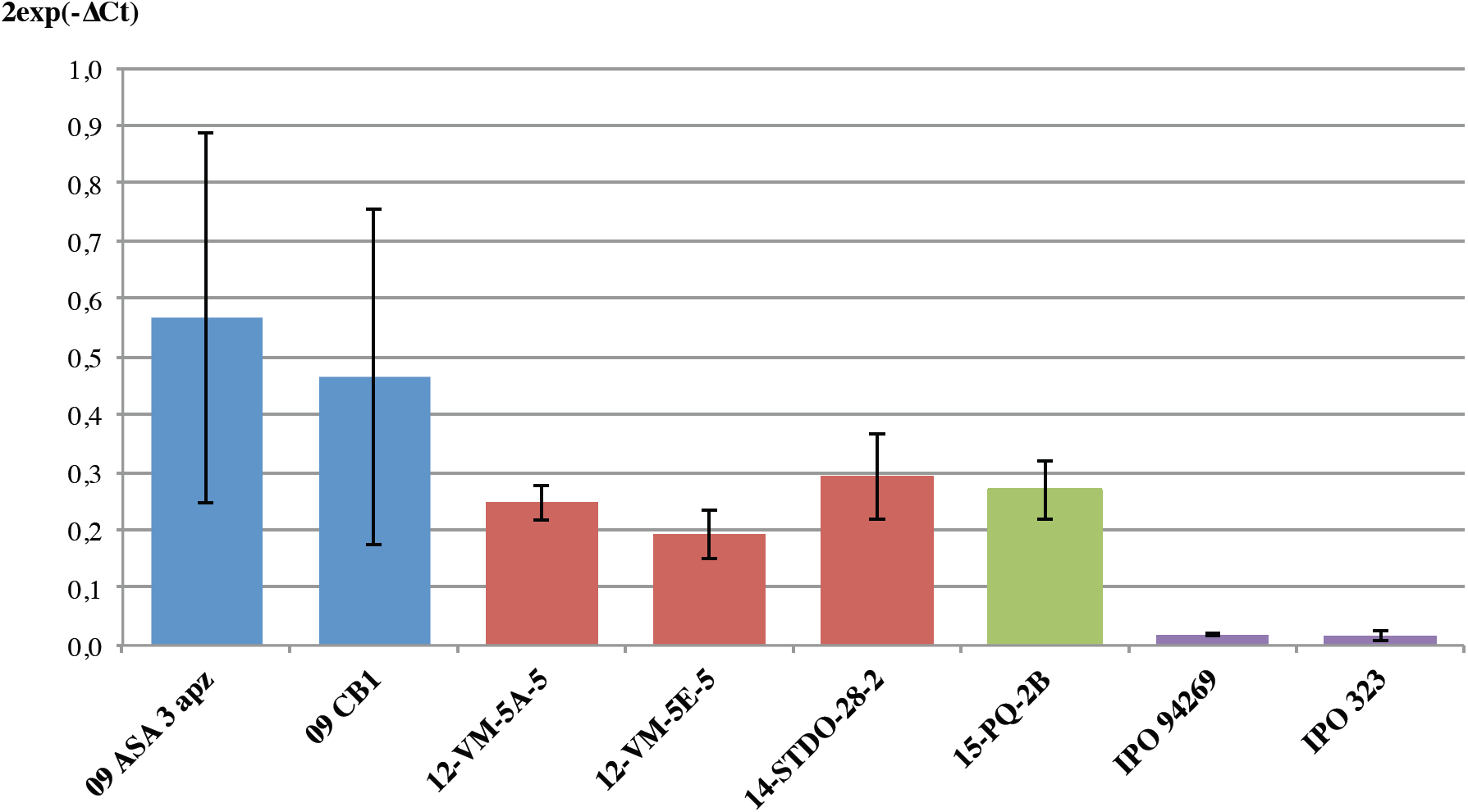
*MFS1*expression in *Z. tritici*field isolates with different promoter genotypes. *MFS1* expression in selected field strains harboring type I-III inserts (blue = type I; red = type II, green = type III, violet = no insert). *MFS1* expression was measured by qRT-PCR relative to three reference genes (*ß-tubulin, eF1a, actin)* in three technical replicates.

### Sequence of the *MFS1* promoter in *Z. tritici* MDR field isolates

We have sequenced the region 500 bp upstream of the *MFS1* start codon in 26 MDR field isolates (Genbank accessions MF623010-MF623033; Fig. S1). This region covers the three types of inserts that localize in a region between 200 and 500 bp upstream of the start codon (Fig. 7D). The integration sites all displayed short repeated sequences (5-10 bp). Type II insert was found in two lengths, 369 bp (type IIa) and in a 30 bp shorter version (type IIb), but otherwise 87% identical. Type III inserts were identical between both strains, 149 bp in length.

**Fig. 7:**
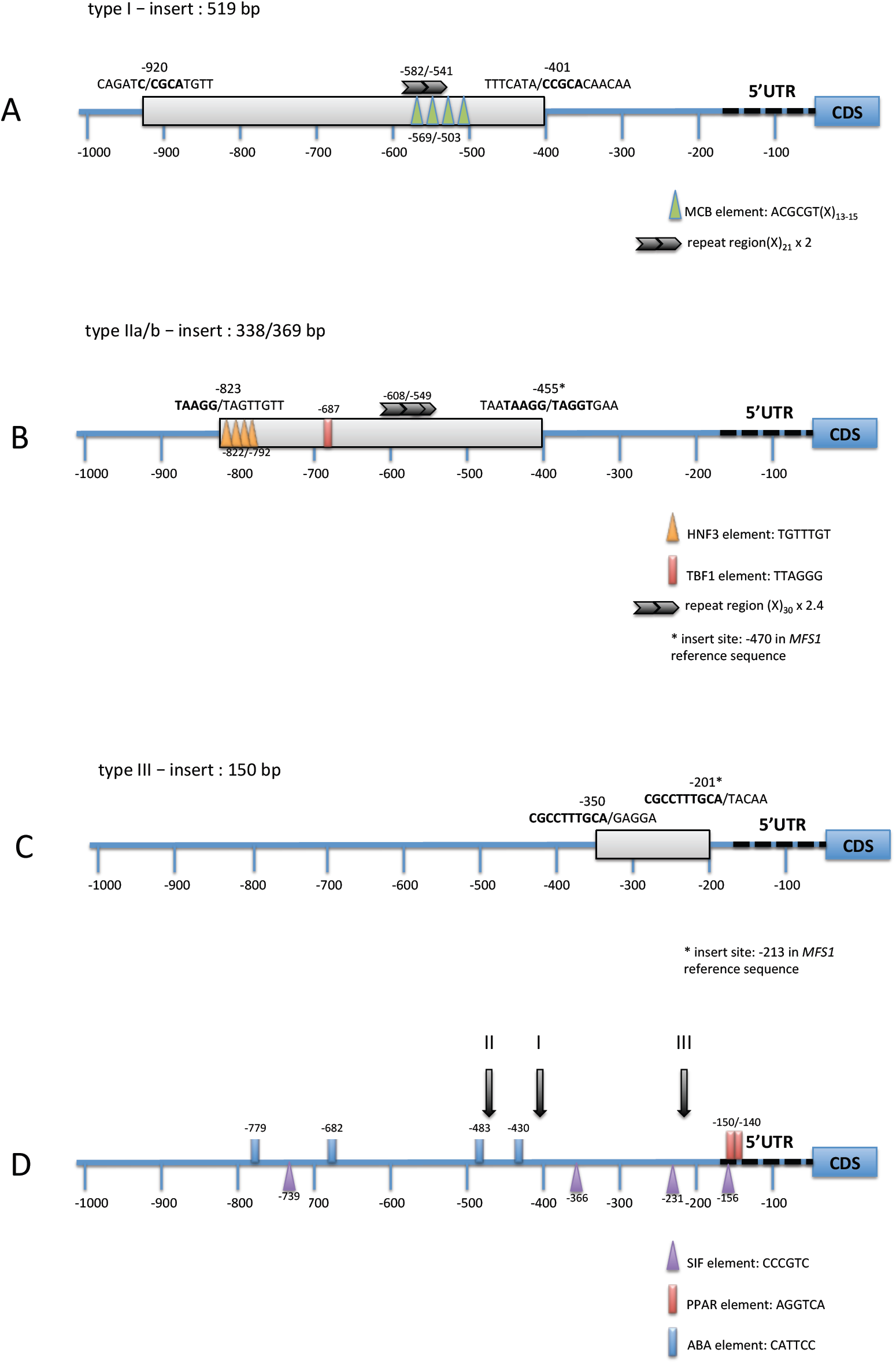
Molecular structure of MFS1 promoter genotypes. The figures indicate the location, length and insert sites of *MFS1* promoter inserts. Repeats between both insertion sites are indicated by bold letters. Consensus sequences of potential regulatory elements identified by TRANSFAC searches (Fogel *et al.*, 2005), are indicated by the colored triangles and boxes. Repeats of longer sequences, found using mreps (Kolpakov *et al.*, 2003), are indicated by the black arrows. For details see text.

As mentioned earlier, the type I insert corresponds to the 519 bp LTR sequence of a Ty1/Copia retrotransposon localized on chromosome 18 (Omrane *et al.*, 2015) in the sequenced strain IPO323 and annotated as still active (Dhillon *et al.*, 2014, Goodwin *et al.*, 2011).

We also screened the sequence of the type II insert against available *Z. tritici* sequences by blastn-searches (Altschul *et al.*, 1997). The query sequence got multiple hits against the IPO 323 sequence (51 hits with >95% identity over >200 bp), had three matches on the genome sequence of JGIBBWB-9N22 and was found 99% identical to the insert found upstream of *CYP51* in some *Z. tritici* DMI resistant isolates, in particular in strain 09-ASA-3apz (Omrane *et al.*, 2015). These results indicate that the type II insert may be a repeated element of the *Z. tritici* genome or part of it.

Searching the sequence of type III insert against the IPO323 genome sequence also gave five hits with >86% identity over more than 110 bp with not annotated nucleotide sequences.

### Identification of putative regulatory elements in the *MFS1* promoter

Since type I insert drives constitutive overexpression of the *MFS1* gene, we suspected this element to contain upstream activating sequences (UAS). UAS may act to recruit transcription activators or co-activators increasing the affinity of the general transcription machinery or through opening the chromatin structure (reviewed in Hahn & Young, 2011) leading to higher transcription levels. Within the 519 bp insert sequence, we searched for known transcription factor binding sites of the TRANSFAC database (Fogel *et al.*, 2005) as well as for larger tandem repeats or palindromic repeats using mreps (Kolpakov *et al.*, 2003). As highlighted in Fig. 7A, we identified four successive consensus sequences of the MCB element (ACGCGT), separated by 13-15 bp respectively in a region 569 to 503 bp upstream of the start codon. This region also overlaps a tandem repeat of 21 bp (−582 to −541). The MCB element (*MluI* cell cycle box, also termed MBP element) is a hexamer sequence regulating the expression of genes involved in G1-phase transition, firstly identified in *S. cerevisiae* (Dirick *et al.*, 1992). The number of MCB repeats correlates with the expression level (Verma *et al.*, 1992). As MCB elements are well conserved in the promoters of G1-phase genes among fungal species (Gasch *et al.*, 2004), one may suspect that the four elements present in the type I insert drive *MFS1* expression in strains harboring this insert.

The same strategy was applied to identify putative UAS in type II and type III inserts. A search against all available consensus site sequences of the TRANSFAC databases lead to multiple hits of more or less randomly distributed sequences (not shown). We therefore limited our search to hexamers and longer sequences, especially those present as direct or inversed repeats. While in the shortest insert (type III) no repeated sequence above this threshold was identified, type II insert harbors different potential regulatory elements as highlighted in Fig. 7B. In a region comprised between −822 and −792 we found four direct repeats of the HNF3 (*Rattus norvegicus*) motif (Cirillo *et al.*, 2002, Johnson *et al.*, 1995). The hexamer TTAGGG corresponding to the *S. cerevisiae* TBF1 element (Brigati *et al.*, 1993) was present at position −687. In addition a 30 bp sequence was repeated 2.4 times in a region between −608 and −549, suggesting that also the type II insert may activate the transcription of the downstream *MFS1* gene.

The original *MFS1* promoter of the sensitive IPO323 strain was analyzed for potential regulatory sequences as described above. Among repeated highly conserved consensus sequences, we identified four repeats of the *Aspergillus nidulans* AbaA binding site CATTCC (Andrianopoulos & Timberlake, 1994), 779 to 430 bp upstream of the start codon, four repeated SIF elements (Bhat *et al.*, 1994, Wagner *et al.*, 1990)(positions −739, −366, −231, −156) and two adjacent PPAR (or PPRE) elements (Blanquart *et al.*, 2003) in the 5’UTR. Some of these elements may be involved in the regulation of *MFS1* transcription under fungicide treatment (Omrane *et al.*, 2015) or under yet unknown conditions.

## Discussion

Increased drug tolerance through increased efflux is a general phenomenon threatening major clinical treatments, anti-cancer, antibacterial and antifungal drugs, respectively (Avner *et al.*, 2012, Wong, 2017, Wu *et al.*, 2014, Paul & Moye-Rowley, 2014). In agriculture, the same phenomenon resulting in MDR has been documented in the last decades with the identification of the involved membrane transporters (Chapeland *et al.*, 1999, Kretschmer *et al.*, 2009, Sang *et al.*, 2015, Leroux *et al.*, 2013, Leroux & Walker, 2013, Omrane *et al.*, 2015). While drug efflux *per se* does not confer high resistance levels, the combination with specific resistance mutations, can strongly impair the efficacy of agro-chemical fungicides. This phenomenon is already being observed in *B. cinerea* (Kretschmer *et al.*, 2009, Leroch *et al.*, 2013, Leroux & Walker, 2013), *S. homeocarpa* (Hulvey *et al.*, 2012, Sang *et al.*, 2015) with several modes of action and, in the case of *Z. tritici*, with DMIs (Leroux & Walker, 2011). The introduction of new modes of action will help dealing with the diseases despite complicated resistance situations, but intelligent treatment strategies must be applied to delay and limit the risk of development and recombination with already fixed resistances.

The mutations responsible for clinical antifungal MDR identified so far, are principally gain-of-function mutations in the transcription factors CaTac1, CaMrr1 (*C. albicans*) and CgPdr1 (*C. glabrata*) regulating the expression of ABC or MFS transporter genes (reviewed in Morschhäuser, 2010, Paul & Moye-Rowley, 2014). In the plant pathogenic fungus *Botrytis cinerea,* similar gain of function mutations in the transcription factor BcMrr1 have been identified in MDR field strains, responsible for the overexpression of the ABC-transporter encoding gene *BcatrB* (Kretschmer *et al.*, 2009, Leroch *et al.*, 2013). In addition, the mutation leading to the overexpression of the MFS transporter encoding gene *BcmfsM2* was identified as a promoter deletion-insertion event; the insert being a retro-element like gene fragment (Kretschmer *et al.*, 2009).

In our previous study, we identified MFS1 as a major player in *Z. tritici* multidrug resistance (Omrane *et al.*, 2015) and had found a retro-transposon relic as an insert in the *MFS1* promoter. In this study we used an unbiased approach by classical and high throughput genetics to identify the MDR-responsible mutations.

We discovered that the *mdr* mutations of both analyzed strains are identical, although their MDR phenotypes differ slightly (Leroux & Walker, 2011), and their drug efflux is affected differently by chemical modulators (Omrane *et al.*, 2015). Therefore one may suspect additional mutations explaining such differences. The observed segregation between sensitive *vs.* MDR strains among the progeny was close to the ratio of 1:1, but the selection of offspring was not completely random. A potential bias can have its origin in the analyzed progeny, as we eliminated 25% of the offspring as mixtures/impure strains. In addition, the phenotypic screen for the MDR phenotype may have been too stringent to identify slight, quantitative differences in resistance to tolnaftate that could have originated from multiple quantitative mutations.

The present study identified three different types of inserts in the *MFS1* promoter region potentially involved in *MFS1* overexpression and, consequently, in MDR. The most frequent insert, type I, is the previously identified relic of a still active Ty1/Copia retrotransposon (Dhillon *et al.*, 2014); the type II insert also resembles a repetitive element, as it was detected at >100 instances in the *Z. tritici* genome sequence. Also the type III insert was found repeated, but less frequently. We have shown that the LTR insert (type I) is responsible for *MFS1* overexpression through classical and reverse genetics. Using a similar gene replacement strategy, we could also confirm the role of inserts II and III in MDR (data not shown).

We addressed the question if the inserts drive *MFS1* expression on their own or if they disrupt transcription repression, through *in silico* analysis of the generated promoter sequences. In the case of the LTR insert, it is highly probable that the insert drives *MFS1* expression on its own, as LTR elements harbor *cis-*regulatory sequences (reviewed by (Chuong *et al.*, 2017, Muszewska *et al.*, 2011)). The *MFS1*-type I LTR element harbors four potential MCB elements known as sequence regulating expression of G1-phase genes in *S. cerevisiae* (Dirick *et al.*, 1992, Moll *et al.*, 1992). As the sequence of this element was found highly conserved upstream of G1-phase genes in fungi (Gasch *et al.*, 2004), these elements probably drive the strong constitutive expression of *MFS1* in type I-MDR strains. Moreover, the regulation of transcription according to cell cycle progression by MCB elements may explain the strong variation observed in the *MFS1* expression levels in type I-MDR strains. Potential UASs were also detected in the type II insert eventually driving *MFS1* constitutive overexpression, while the type III insert seems devoid of novel regulatory elements. In sensitive *Z. tritici* strains, *MFS1* expression is induced after fungicide challenge (Omrane *et al.*, 2015), while all insert leads to high basal expression. These results suggest that the LTR insert drives expression on its own while abolishing the induction under fungicide challenge (Omrane *et al.*, 2015). The function of type II insert might be similar (induction instead of release of repression), while in the case of type III insert, its position close to the 5’ UTR and the absence of known UASs, we may suspect that this element rather releases the inhibition of *MFS1* transcription. To fully understand the regulation of MFS1 expression, to discriminate between both hypotheses and the role of the potential regulatory elements in *MFS1* transcription regulation remains to be established through promoter fusion experiments. In addition the transcription regulators involved in *MFS1* regulation in response to drug challenging remain to be identified in *Z. tritici*.

Finally, our analysis of *Z. tritici* field strains revealed three to four different insertion events into the *MFS1* promoter with strong impact on the fungicide sensitivity phenotype. Mobile elements mediating fungicide resistance through target gene overexpression is a common mechanism among phytopathogenic fungi. Especially in the *CYP51* promoters of various fungal species repeated elements or reminiscence of them were found, *e.g., Penicillium digitatum* (Ghosoph *et al.*, 2007, Sun *et al.*, 2013), *Blumeria jaapii* (Ma *et al.*, 2006), *Monilinia fructicola* (Luo & Schnabel, 2008), *Venturia inequalis* (Schnabel & Jones, 2001, Villani *et al.*, 2016), and *Z. tritici* (Cools *et al.*, 2012). Transposable elements (TEs) are known to modify genome structure, gene functions and phenotypes (Castanera *et al.*, 2016, Chuong *et al.*, 2017, Hirsch & Springer, 2016, Amselem *et al.*, 2015). In filamentous fungi, the genome content in TEs can be highly variable even between closely related species (Muszewska *et al.*, 2011, Grandaubert *et al.*, 2014). TEs are suspected to contribute to host adaptation, *e.g.,* in *Magnaporthe oryzae* (Yoshida *et al.*, 2016), to speciation and to higher adaptive capacity to selective pressure (including pathogenicity evolution) by generating genomic rearrangements (Grandaubert *et al.*, 2014, Hartmann & Croll, 2017). It was recently established that >17% of the *Z. tritici* genome sequence was repetitive, out of which 70% corresponding to retro-transposable elements (Dhillon *et al.*, 2014, Grandaubert *et al.*, 2015).

Among the LTR-transposons identified by Dhillon et al. (Dhillon *et al.*, 2014), one (family 18) showed minimal evidence of RIP and was therefore supposed to be still active. Indeed, the LTR-sequence corresponding to the MFS1-type I insert, is 100% identical to this family 18 LTR. We may therefore suspect that this LTR insertion is due to a recent retro-transposition event. According to their data, Grandaubert and co-workers supposed that also class II DNA transposons are still active in *Z. tritici* (Grandaubert *et al.*, 2015). They might be at the origin of the other two insertion events.

Little is known about the influence of fungicide exposure on (retro-) transposon mobilization. Chen and colleagues observed increased mobilization of the transposable element *Mftc1* in *M. fructicola* after *in vitro* exposure to sub-lethal fungicide concentrations, although not affecting fungicide sensitivity (Chen *et al.*, 2015).

The question remains if the *MFS1* promoter is prone to insertion events relatively more frequently than other genomic *loci* or if this is only due to fungicide selection pressure. The available and forthcoming *Z. tritici* whole genome sequences and associated transcriptomic data will shade light on the influence of transposable elements on genome structure and global gene expression (Grandaubert *et al.*, 2015, Zhong *et al.*, 2017, Hartmann *et al.*, 2017), but also on their evolution. Population genomics and experimental evolution may also help evaluating the impact of fungicide pressure on transposon mobilization.

## Acknowledgements

The authors are grateful to Arvalis, Institut du Végétal, to BASF, Bayer, DuPont de Nemours and to Syngenta for financial support. Part of the study was financed by Ministère de l’environnement, du développement durable et de l’écologie (MEDDE) through the Ecophyto II program.

